# The Electrophysiological Signature of Mind Wandering

**DOI:** 10.1101/819805

**Authors:** Stefan Arnau, Christoph Löffler, Jan Rummel, Dirk Hagemann, Edmund Wascher, Anna-Lena Schubert

**Affiliations:** Leibniz Research Centre for Working Environment and Human Factors Dortmund (IfADo), Dortmund, Germany; Heidelberg University, Institute of Psychology, Heidelberg, Germany

**Keywords:** Mind wandering, EEG, time frequency analysis, alpha, resource allocation

## Abstract

Mind wandering during ongoing tasks can impede task performance and increase the risk of failure in laboratory as well as in daily-life tasks and work environments. Neurocognitive measures like the electroencephalography (EEG) offer the opportunity to assess mind wandering non-invasively without interfering with the primary task. However, the literature on electrophysiological correlates of mind wandering is rather inconsistent. The present study aims towards clarifying this picture by breaking down the temporal dynamics of mind-wandering encounters using a cluster-based permutation approach. Participants performed a switching task during which mind wandering was occasionally assessed via thought probes applied after trial completion at random time points. In line with previous studies, response accuracy was reduced during mind wandering. Moreover, alpha power during the inter-trial interval was significantly increased on those trials on which participants reported that they had been mind-wandering. This spatially widely distributed effect is theoretically well in line with recent findings linking an increased alpha power to an internally oriented state of attention. Measurements of alpha power may therefore be used to detect mind wandering online during critical tasks in traffic and industry in order to prevent failures.

## Introduction

Mind wandering, that is the experience of one’s attention drifting away from the external environment towards inner thoughts and feelings, is a frequent phenomenon. Mind wandering is as an umbrella term for divergent states of inattention (Seli et al., 2018a) but is often conceptualized in a more specific manner, namely as a redirection of attention away from a currently ongoing task (Mrazek et al., 2013). In this paper, we use the term mind wandering in this more specific sense, as a reference to task-unrelated thoughts that occur while one is engaged in some ongoing task.

Indeed, it seems that during many mundane tasks – even when writing a manuscript on mind wandering – we often find ourselves preoccupied with thoughts totally unrelated to our tasks at hand. In one large-scale ambulatory assessment study deploying experience sampling during the daily lives, mind wandering was reported during 46.9% of all daily activities (Killingsworth & Gilbert, 2010). Although the prevalence of mind wandering was found to be lower in some more recent studies (i.e., around 30%; Kane et al., 2017), these findings render mind wandering a relevant factor to deal with when investigating cognitive performance. Decrements in performance due to mind wandering have been observed in sustained attention (Allan Cheyne et al.,2009; Denkova et al., 2018), memory (Risko et al., 2012; Smallwood et al., 2003), working memory capacity (Kane & McVay, 2012; Mrazek et al., 2012), reading (Franklin et al., 2011; Schad et al., 2012), simulated driving (Lemercier et al., 2014; Zhang & Kumada, 2018), and random number generation (Taesdale et al., 1995). In studies aiming towards a higher ecological validity, mind wandering was also found to impair real-world driving performance (e.g. Burdett et al., 2019; Galéra et al., 2012; Qu et al., 2015) as well as daily life performance (McVay et al., 2009).

One limiting factor for the investigation of mind wandering is its assessment. Most research on mind wandering relies on participants self-reported mental states. That is, participants either need to catch themselves when they are off task by pressing a designated key (self-caught method) or they are randomly asked from time to time during an ongoing task whether they had just been on task or off task before the thought probe occurred (probe-caught method). Although both methods have been used very successfully for studying the wandering mind (Smallwood & Schooler, 2015), they also impose some methodological limitations (see Weinstein (2018) for a review). To capture all mind wandering instances, the self-caught method would require participants to be fully aware of their own mental states all the time while performing a task, which is often not the case (Smallwood et al., 2007). The probe-caught method captures mind wandering with and without awareness but requires participants to temporarily interrupt their primary task to report on their thoughts. Although mind wandering probes do not interfere much with task performance, at least during simple cognitive tasks (Wiemers & Redick, 2018), they still interrupt the ongoing task and thus alternative indicators of mind wandering are needed (cf. Steindorf & Rummel, 2019). Different mind wandering indicators have been suggested, such as changes in response time variabilities (McVay & Kane, 2012), changes in pupil dilation (Unsworth & Robison, 2018), or changes in neural activity (Christoff et al., 2009). The last method is particularly promising, because identifying a neuro-cognitive signature of mind wandering would not only allow to assess mind wandering without thought probes but also to better understand the temporal dynamics of mind wandering episodes when using methods with high temporal resolutions such as EEG. The present study thus aims to investigate the neuro-cognitive signature of mind wandering by means of the EEG.

Some previous studies investigated the relationship between mind wandering and EEG activity. In most studies, Event Related Potentials (ERP) responses were found to be decreased during mind wandering episodes. In a study comprising simulated driving as well as cognitive tasks, Baldwin and colleagues (2017) found reduced P3a amplitudes in mind wandering episodes at frontal and central recording sites. Barron and coworkers (2011) used a retrospective self-report measure to compare the electrophysiological data of a high versus a low mind wandering group in an oddball paradigm. They observed reduced ERP responses to targets (P3b) as well as to standard (P3a) in the high mind wandering group. Decreased P3 components during mind wandering were also observed by Kam & Handy (2013). The P3 amplitude has repeatedly been associated with the allocation of cognitive resources (Allison & Polich, 2008; Kok, 2001). Reduced ERP amplitudes during mind wandering have also been observed for early sensory components such as the visual P1 (Baird et al., 2014; Kam & Handy, 2013) and the auditory P2 (Braboszcz & Delorme, 2011).

When considering EEG oscillatory activity, theta and alpha band dynamics might be of particular interest in the context of mind wandering. Theta, especially event-related frontal-midline theta activity has been associated with the exertion of cognitive control (Cavanagh et al., 2012; Cavanagh & Frank, 2014). Cognitive control allocated to a task should be decreased in mind wandering situation, as attentional resources are drawn away from the primary task (Mrazek et al., 2013). Alpha activity, on the other hand, was found to be suppressed by sensory stimulation (Thut et al., 2006) and increased in periods without stimulation (Carp & Compton, 2009; Compton et al., 2011). Alpha power seems to reflect an internally oriented attentional state (Hanslmayr et al., 2011). Consequently, it was found to be increased during mental imagery (Cooper et al., 2003) and has been associated with default mode network activity (Knyazev et al., 2011; Mo et al., 2013). An internal focus of attention and mental imagery are relevant aspects in mind wandering situations, which renders alpha power a promising variable to reflect certain aspects of mind wandering in the EEG.

The findings of previous research with respect to the representation of mind wandering in time frequency space are rather inconclusive. Atchley and colleagues (2017) used a Stroop task to assess mind wandering by defining segments preceding error-trials as mind wandering episodes. They found the spectral power in the theta and alpha band to be decreased compared to segments preceding correct trials. Baldwin et al. (2017), however, found an increase of alpha power during mind wandering at parietal electrodes using the probe-caught assessment method during simulated driving as well as during a vigilance task. Similar results could be observed by Compton et al. (2019), who also found increased alpha power in episodes preceding reports of mind wandering during a Stroop task. Baird and colleagues (2014) also used the probe-caught method to investigate mind wandering during a vigilance task. They observed a larger event-related reduction of alpha power in the Event Related Spectral Perturbation (ERSP) of the EEG during mind wandering episodes at frontal and parietal leads as compared to on-task trials. Braboszcz and Delorme (2011) combined breath counting as the primary task with the self-caught method to assess mind wandering. Braboszcz and Delorme (2011) compared EEG segments that preceded button presses indicating self-caught mind wandering with episodes after the button press. Using a cluster-based permutation approach (Maris & Oostenveld, 2007), Braboszcz and Delorme (2011) found significantly increased delta and theta power in the pre-button-press segment as compared to the post-button-press segment at all electrodes. The result pattern was opposite for the alpha and beta band. For the alpha band, however, the lower power in the pre-compared to the post-button-press segment was only significant at occipital electrodes. Other studies investigated spectral power ratios in order to identify reliable correlates of mind wandering in the EEG. Applying the self-caught method, van Son et al. (2019) found increased *theta*/*beta* ratio at frontal electrodes during mind wandering. Similar findings have been reported for the *beta*/*alpha* ratio and the *beta*/(*alpha* + *theta*) ratio (Cunningham et al. 2000).

The inconsistency in previous findings on the electrophysiological representation of mind wandering is somewhat unsatisfactory and most likely due to the large variety of methods used. The choice of signal processing methods may affect the results drastically. The choice of electrodes used, whether absolute or relative power measures are used, and the kind of baseline correction normalization may affect the outcome substantially. The assessment of mind wandering appears to be crucial in particular as the findings of recent research with respect to the alpha band seem be the opposite for the probe-caught compared to the self-caught method. It might be questionable, whether the pre-response period in the self-caught approach reflects mind-wandering, the cognitive process of getting aware of mind-wandering, response preparation, or a mixture of these processes. Finally, the context or tasks in which mind wandering is assessed may play an important role for the observed inconsistencies in the cognitive signature. Mind wandering is more prevalent and interferes less with task performance in low as compared to high demand tasks (Robison & Unsworth, 2018; Rummel & Boywitt, 2014). Mind wandering is also more prevalent in practiced versus non-practiced tasks (Cunningham et al., 2000; Giambra, 1995) in automated environments (Gouraud et al., 2018), that is situations in which less attentional control is needed, and when probed less frequently (Schubert et al., 2019a; Seli et al., 2013).

The present study aims towards obtaining a clear electrophysiological signature of mind wandering, as far as possible. To this end, participants performed a switching task. The assessment of mind wandering was done by applying the probe-caught method in which the participants were intermittently asked whether they experience mind wandering at that respective moment or not. Trials in which mind wandering was reported as well as the two preceding trials were defined as mind wandering trials. Other trials were defined as on-task trials. In order not to focus on specific electrodes and also to address the problem of multiple comparisons, a cluster-based permutation approach was chosen to statistically test for differences between mind wandering and on-task episodes (Maris & Oostenveld, 2007). As we also wanted to avoid a priori assumptions about the temporal structure of effects of mind wandering in relation to the primary task, we analyzed segments covering task-related processing as well as the inter-trial interval. We chose a decibel normalization approach using the frequency-specific power averaged across all conditions, that is mind wandering and on-task episodes, in a pre-stimulus interval as baseline. In doing so, systematic mind-wandering related variance during the baseline period would not be reflected to time-frequency points outside the baseline. The switching task used as the ongoing task in the present study has the additional advantage that it provides contexts in which task demands are higher (i.e., on switching trials) and contexts in which task demands are lower (i.e., on repeat trials). Beside the factor mind wandering versus on-task trials, we thus also tested for effects of switch versus repeat trials, as well as for the interaction.

## Methods

### Sample

For the experiment, 100 participants were recruited via local newspaper advertisement, announcements on social media platforms, as well as via the distribution of flyers in Heidelberg. The inclusion criteria were an age between 18 and 60 years, normal or corrected to normal vision, and not having a history of mental illness. Two participants had to be excluded for not completing the experiment. For the present study, we only considered those participants that reported to have experienced mind wandering at least at 30 occasions during the experimental task. This reduced our sample size to *N* = 32 (19 females) with a mean age of 29.88 years (*SD* = 11.9). All participants signed an informed consent before participating in the experiment. The study was approved by the ethics committee of the Heidelberg University and all methods used are in accordance with the declaration of Helsinki.

### Procedure

The experiment started immediately after participants completed an intelligence test (data reported in Schubert et al., 2019b) and after the preparation of a 32 electrode EEG montage. Stimuli were presented using MATLAB 2017b (The MathWorks Inc., Natick, Massachusetts) against a black background on a 21.3” monitor (EIZO FlexScan S2100) with a resolution of 1600 × 1200 pixels and a refresh rate of 60 Hz. The participants performed in a shifting task (Sudevan & Taylor, 1987) during which digits ranging from 1 to 4 and 6 to 9 were presented at the central position on the screen that subtended a visual angle of 0.6° at a viewing distance of approximately 74 cm. Depending on the color of the digit, participants had either to decide whether the presented number was odd or even, or if the number was less or more than 5. The colors indicating the task were red (#F00000) for the less/more task and green (#00F000) for the odd/even task. After 40 practice trials which included feedback, each participant completed 640 trials without feedback that were divided into ten blocks with 64 trials each. Both tasks as well as whether the task changed from one trial to the next (switch-trial) or stayed the same (repeat-trial) were equally likely for each trial. Responses had to be given by pressing the D and L buttons on a computer-keyboard with the left and right index fingers. In order to prevent systematic variance due to anticipation, the temporal structure of the trials was jittered. Each trial started with the presentation of a fixation cross in grey (#787878, 0.6° visual angle) for 512 to 768 ms, followed by a blank screen for 1024 to 1278 ms. Subsequently the imperative stimulus, that is the colored digit, was presented until 1024 to 1278 ms after the participants’ response. The following inter-trial interval was 1000 to 1500 ms long. The temporal structure of the trials is depicted in Figure 1. After a trial, there was a random chance that the participants were asked *“Where have you just been with your thoughts?”*. The participants were asked to respond with the left arrow button if they were focused on the task and to answer with the right arrow button when they experienced mind wandering. The minimum distance between these experience probes was 5 trials and the maximum distance was 10 trials. Trials for which the participants reported mind wandering as well as the two preceding trials were defined as mind wandering trials, whereas other trials were defined as on-task trials.

**Figure 1:**
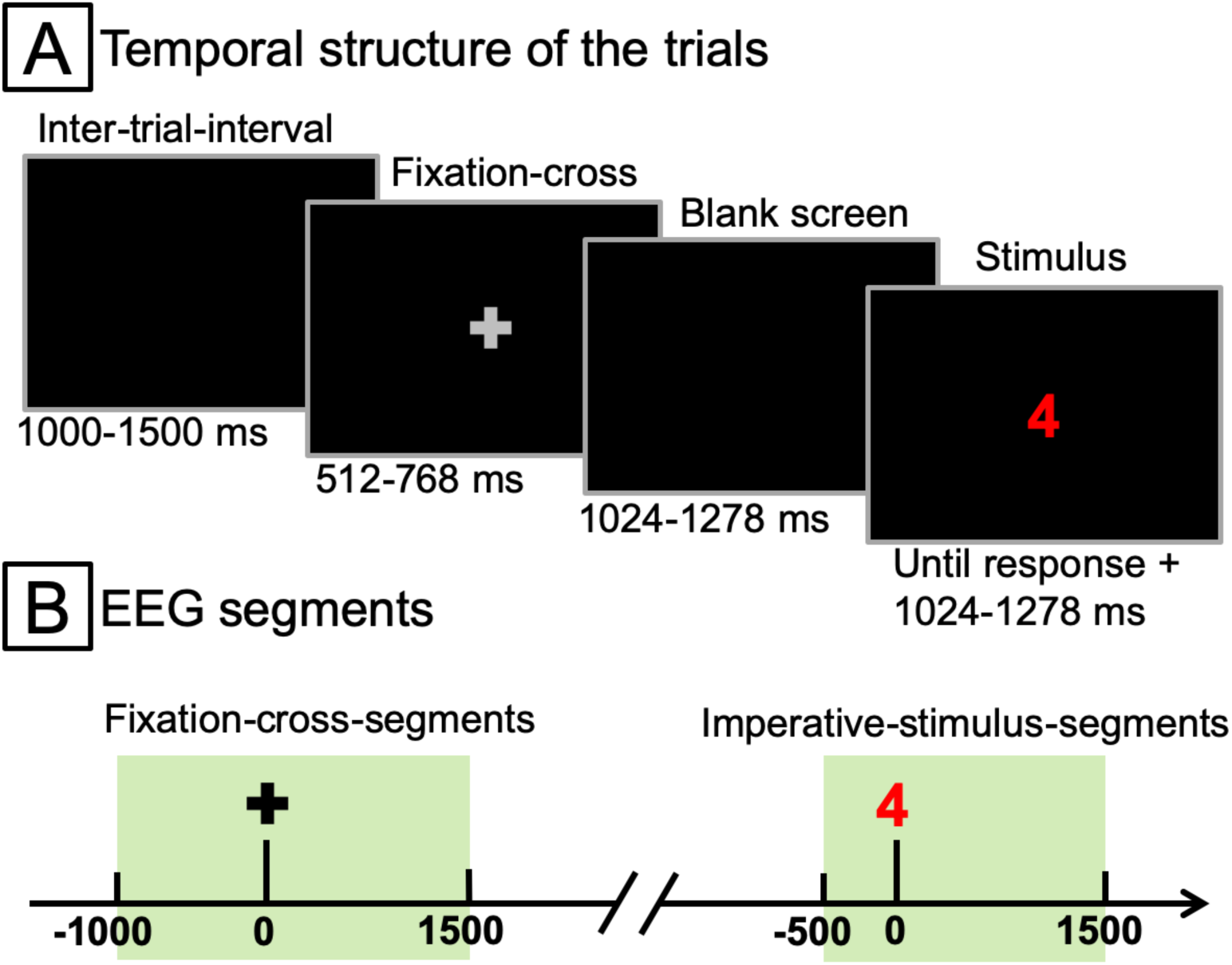
This figure depicts the temporal structure of the trials [A] as well as the position of the fixation-cross-locked and stimulus-locked segments used for the EEG analysis within the trials [B].

### EEG recording

The EEG recording took place in a dimly lit and sound-attenuated cabin. A montage of 32 Ag/AgCl EEG electrodes was used, which were equidistantly distributed on the scalp (Equidistant Montage No. 7, Easycap GmBH, Herrsching, Germany). The ground electrode was placed at AFz and Cz was used as online reference. The EEG data were sampled at 1000 Hz using BrainAmp amplifier (Brain Products, Munich, Germany). A 0.1 Hz hardware hi-pass filter was used and impedances were constantly kept below 5 kΩ.

### EEG preprocessing

Signal processing and analysis of EEG data was performed in Matlab 2018b (The MathWorks Inc., Natick, Massachusetts) using custom scripts incorporating functions of the EEGLab (Delorme and Makeig, 2004) and fieldtrip (Oostenveld et al., 2011) toolboxes. The data were band-pass filtered at 1 to 30 Hz and corrupted channels were identified and removed based on kurtosis and probability criteria. On average 0.97 channels (*SD* = 0.93) were removed. Subsequently, data were re-referenced to common average reference, resampled at 200 Hz, and segmented into epochs ranging from −2000 ms to 2000 ms relative to the onset of a fixation cross or imperative stimuli. Fixation cross and imperative stimulus events were segmented separately due to the temporal jitter in the inter stimulus interval as well as of the fixation cross presentation time (see Figure 1[A]). After the automatic detection and removal of epochs containing artifacts, an independent component analysis was performed. Independent components (ICs) representing artifacts were identified and removed using ICLabel (Pion-Tonachini et al., 2019) by retaining only ICs which were labeled in the *Brain* IC-category with a probability of at least 0.5. On average, 11.50 ICs (*SD* = 3.52) were excluded. Again, corrupt epochs were rejected automatically and all fixation cross and imperative stimulus segments without the respective counterpart were removed as well. On average, 235.88 (*SD* = 17.70) fixation-cross-locked and imperative-stimulus-locked segments entered the analysis of behavioral and electrophysiological data.

### Time frequency decomposition

A time frequency decomposition of the data was performed by convolving the electrophysiological data with complex Morlet wavelets defined as complex sine waves tapered by a Gaussian. A set of 50 wavelets was used with frequencies ranging from 2 to 30 Hz in logarithmically spaced steps. The widths of the corresponding tapering Gaussians were defined in a way that the resulting wavelets had a temporal resolution ranging from 600 to 50 ms at full-width at half-maximum (FWHM; Cohen, 2018), which corresponds to a FWHM ranging from 1.25 to 17.25 Hz in the frequency domain. Power estimates were extracted squaring the absolute values of the complex convolution result. For the group-level analysis, data were subsequently decibel normalized in time-frequency space relative to a baseline ranging from − 500 to −200 ms before the onset of a fixation-cross or an imperative stimulus. It is important to note that the baseline was calculated based on all trials. In contrast to applying a baseline normalization specifically for each condition, or even for each trial, variance related to the experimental condition thus remains present in the baseline period and may be observed there. When applying a baseline normalization of the data in a condition-specific manner, however, systematic variance that may be present in the baseline period would be shifted to extra-baseline periods. In order to avoid temporal overlap (c.f. Figure 1) and also to remove edge artifacts, fixation-cross segments were pruned to −1000 to 1500 ms and imperative stimulus segments were pruned to −500 to 1500 ms to obtain the final ERSPs.

### Statistics

In order to test for the effect of trial type (repeat vs. switch) as well as of reported mind wandering (on-task vs. mind wandering), linear mixed models were fitted to the behavioral data on single trial level. Response speed and response accuracy entered the respective model as dependent variables and the experimental factors trial type and reported mind wandering entered the model as fixed effects. A random intercept was modeled for each participant (dv ∼ trial type * mind wandering + (1|subject)).

In order to test for significant effects in the electrophysiological measures in time × frequency × sensor space, cluster-based permutation tests were performed (Maris & Oostenveld, 2007). This approach has the advantage of controlling for Type I error rates, which is crucial for dealing with multiple comparisons in high-dimensional data. For each data point in time × frequency × sensor space, *t*-statistics were computed and a clustering algorithm identified clusters of neighboring data points associated with a *t*-value corresponding to *p* < 0.1. The test statistic for each identified cluster was computed as the summed *t-*values of all data-points included. Type I error was controlled for by evaluating this test statistic under a H0 distribution of maximum cluster-level statistics determined in a randomization procedure with 1000 iterations. In each of these iterations the maximum cluster statistic was determined based on data with randomized factor level assignments. Subsequently, the actually observed test statistics were compared against this H0 distribution in a two-sided test. Clusters with p < 0.05 were regarded as significant. Overall, three cluster-based permutation tests were performed. In order to test for significant effects of the trial sequence, the assignment of the data to repeat and switch trials was permuted in the randomization procedure. For the test for effects of mind wandering the randomization procedure utilized reported mind wandering. In order to test the interaction, the differences of repeat and switch trials were computed for each mind wandering level and the result entered the same procedure as the main effects. The tests for effects of trial sequence and the interaction were only performed on segments locked to the imperative stimulus, since the fixation cross did not contain any information about the subsequent task. The test for effects of mind wandering, however, was performed on fixation cross locked segments as well, as we assumed mind wandering to occur during the inter-trial-interval as well. All effect sizes were estimated as bias-corrected partial η^2^ (Mordkoff, 2019; subsequently referred to as η^2^) and classified as small, medium or large according to conventions of Cohen (1992).

## Results

### Behavior

The behavioral data are depicted in Figure 2 and the corresponding test statistics are listed in Table 1. The statistical analysis revealed that the participants responded significantly faster in repeat trials as compared to switch trials (*M* = 843.11 ms, *SD* = 119.76 versus *M* = 910.64 ms, *SD* = 128.32) with *p* < .001. No significant effect could be observed for the factor mind wandering (*M* = 881.82 ms, *SD* = 128.17 for trials without mind wandering and *M* = 871.31 ms, *SD* = 116.90 for trials with mind wandering), neither for the interaction. Response accuracy was not significantly affected by the trial type (*M* = 96.63 %, *SD* = 2.73 in repeat trials versus *M* = 95.76 %, *SD* = 3.09 in switch trials), but was significantly higher in trials in which participants did not report mind wandering than in trials in which participants reported mind wandering (*M* = 97.15 %, *SD* = 3.41 versus *M* = 94.74 %, *SD* = 3.22). The interaction was not significant.

**Table 1:** For response time and accuracy, this table shows *t*-statistics (*t*) with the corresponding effect sizes (η^2^), regression coefficients (β) and the 95% confidence interval (CI) for the fixed effects mind wandering (MW), trial sequence (SEQ, that is switch vs. repeat trials), and their interaction. Test statistics corresponding to *p* < .05, or *p* < .001 are marked with one and two asterisks, respectively.

|  | response time |  |  | accuracy |  |  |
| --- | --- | --- | --- | --- | --- | --- |
| | $t(\eta^2)$ | $\beta$ | 95% CI | $t(\eta^2)$ | $\beta$ | 95% CI |
| **MW** | -1.29(0.02) | -11.28 | -28.46, 5.91 | -2.94*(0.19) | -0.019 | -0.032, -0.006 |
| **SEQ** | -9.43**(0.73) | 71.92 | 56.96, 86.87 | -0.85(-0.01) | -0.005 | -0.016, 0.006 |
| **MW×SEQ** | -0.64(-0.02) | -7.77 | -31.56, 16.01 | -1.03(0) | -0.009 | -0.027, 0.008 |

**Figure 2:**
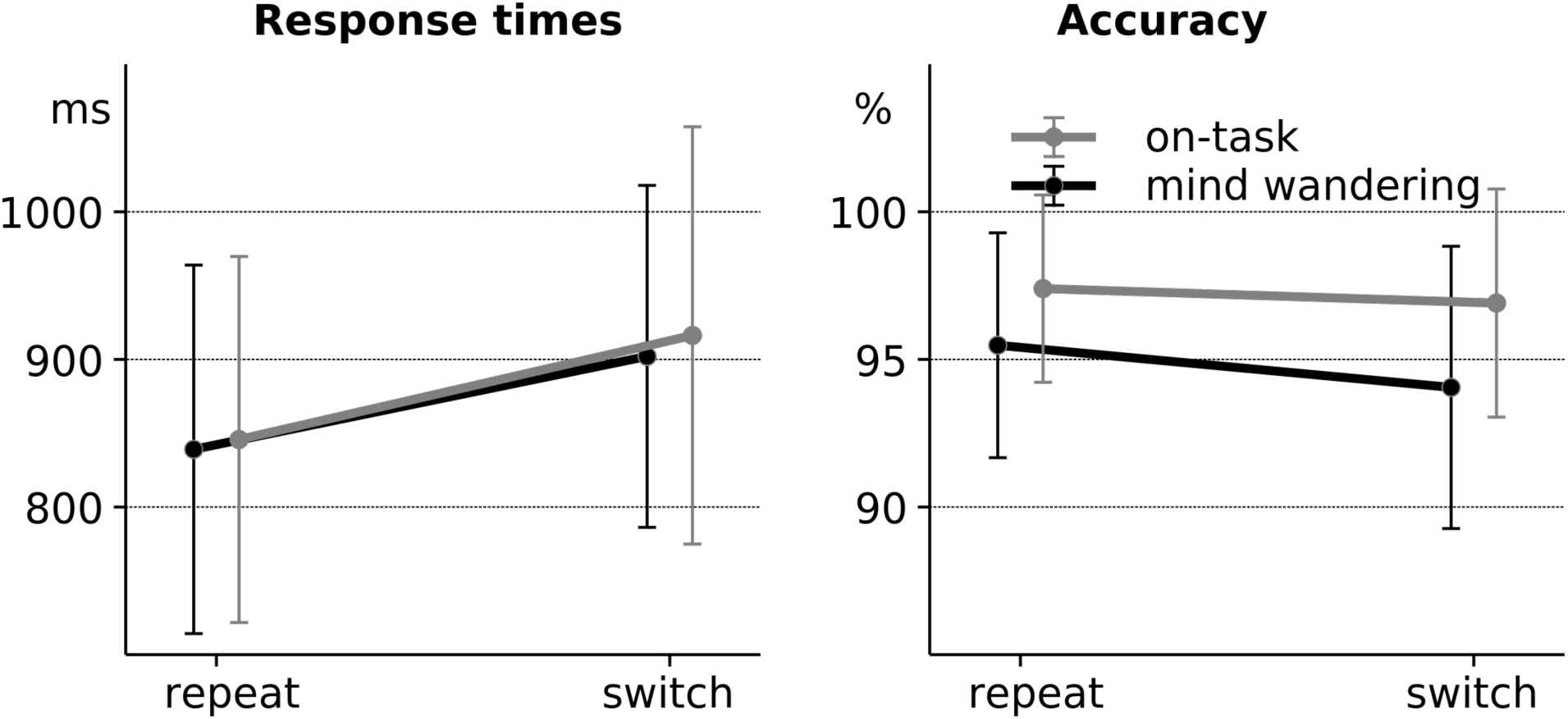
This figure depicts the behavioral measures response time (left panel) and accuracy (right panel). The error bars are representing the standard deviations.

### Time-frequency-decomposition of EEG data

The cluster-based permutation test in time × frequency × sensor space for significant differences in trials with mind wandering versus trials without mind wandering revealed 4 significant clusters, illustrated in Figure 3. All of these 4 clusters had negative sums on *t*-values, indicating a significantly greater spectral power relative to the baseline in trials with reported mind wandering as compared to trials without mind wandering. Cluster 1 is located in the alpha and theta band. It comprises the inter-trial-interval and lasts until approximately 700 ms after the fixation cross onset. The effect size is largest in the alpha band during the inter-trial-interval (see Figure 3 [C]). In sensor space, Cluster 1 is significant at all recording sites and exhibits the largest effect sizes at lateralized central and parietal electrodes. Cluster 2 and Cluster 3 are located in the alpha and theta band as well. These clusters comprise the time from approximately 1000 ms after the onset of the fixation cross (Cluster 2 in the segments locked to the fixation cross) until the onset of the imperative stimulus (Cluster 3 in the segments locked to the imperative stimulus). Cluster 2 and Cluster 3 show the largest effect sizes in the theta band and at posterior leads, although the effect is significant at a large number of electrodes (see Figure 3 [D]). Cluster 4 is located in the alpha, theta, and delta band. It starts approximately 500 ms after the onset of the imperative stimulus and ranges into the inter-trial-interval. Like Cluster 1, it is significant at all recording sites and shows the largest effect sizes in the alpha band (see Figure 3 [C]) and at central and parietal electrodes (see Figure 3 [D]).

**Figure 3:**
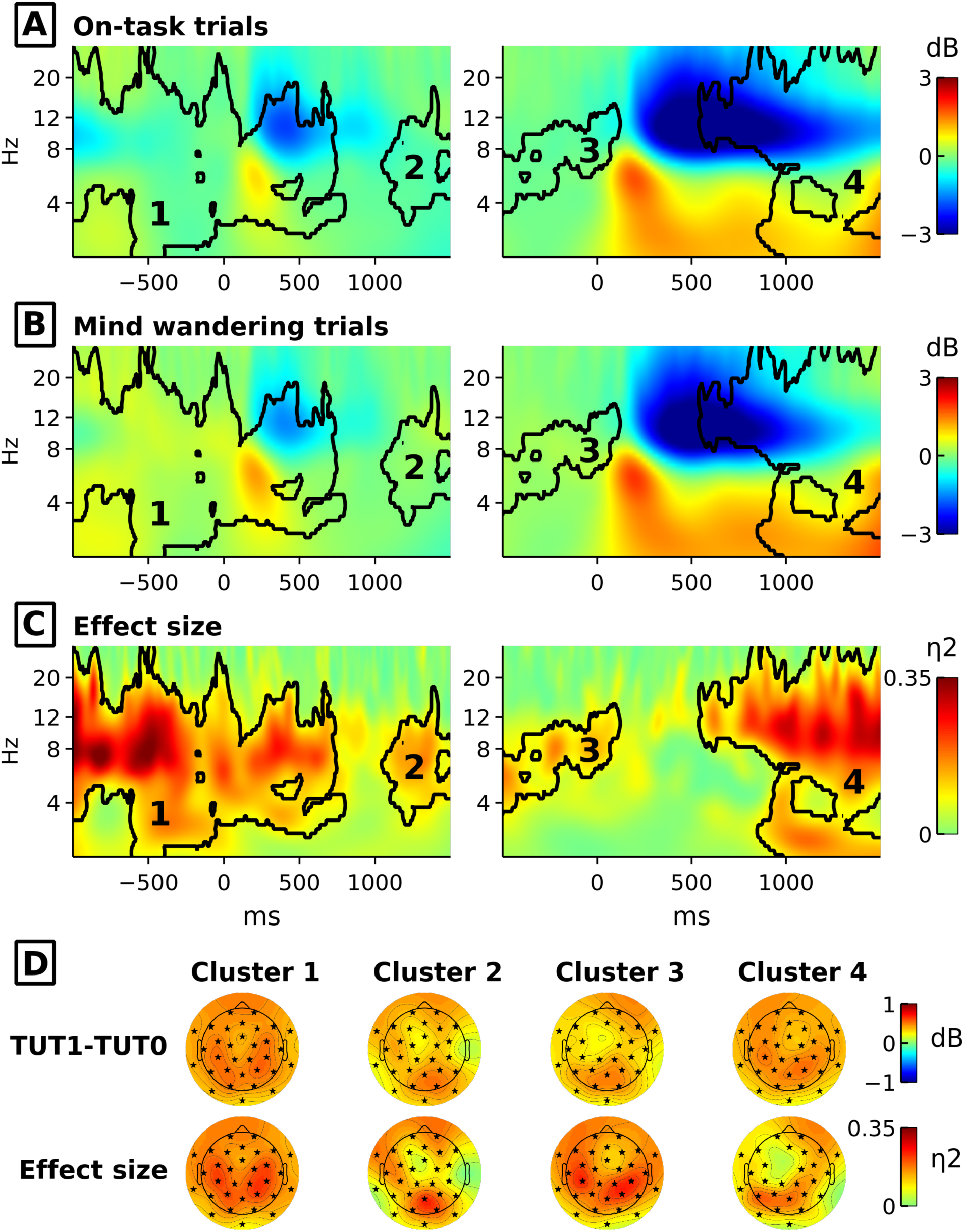
The figure illustrates the four significant clusters for effects of mind wandering. The ERSPs averaged across all channels for on-task trials are depicted at [A] and for mind wandering trials at [B]. Panel [C] illustrates the corresponding effect sizes. The left column of [A], [B], and [C] represent EEG segments locked to the fixation cross, the right side the segments locked to the imperative stimulus. The black contour lines indicate significant clusters and the black numbers represent the cluster number. Panel [D] depicts the topographies for the clusters with the power difference between mind wandering and on-task trials illustrated in the upper row and the corresponding effect sizes illustrated in the lower row. The black asterisks indicate channels with significant effects of the respective cluster.

The cluster-based permutation test for effects of trial type revealed one significant cluster illustrated in figure 4. This cluster, Cluster 5, is positive, thus indicating a greater spectral power relative to the baseline in repeat trials compared to switch trials. Cluster 5 is situated in the alpha and lower beta range and significant at a large number of electrodes (see Figure 4 [D]). The largest effect sizes of Cluster 5 are in the alpha range (see Figure 4 [C]) at frontal and posterior recording sites (see Figure 4 [D]). The cluster-based permutation test for the interaction of the factors trial sequence and mind wandering revealed no significant results.

**Figure 4:**
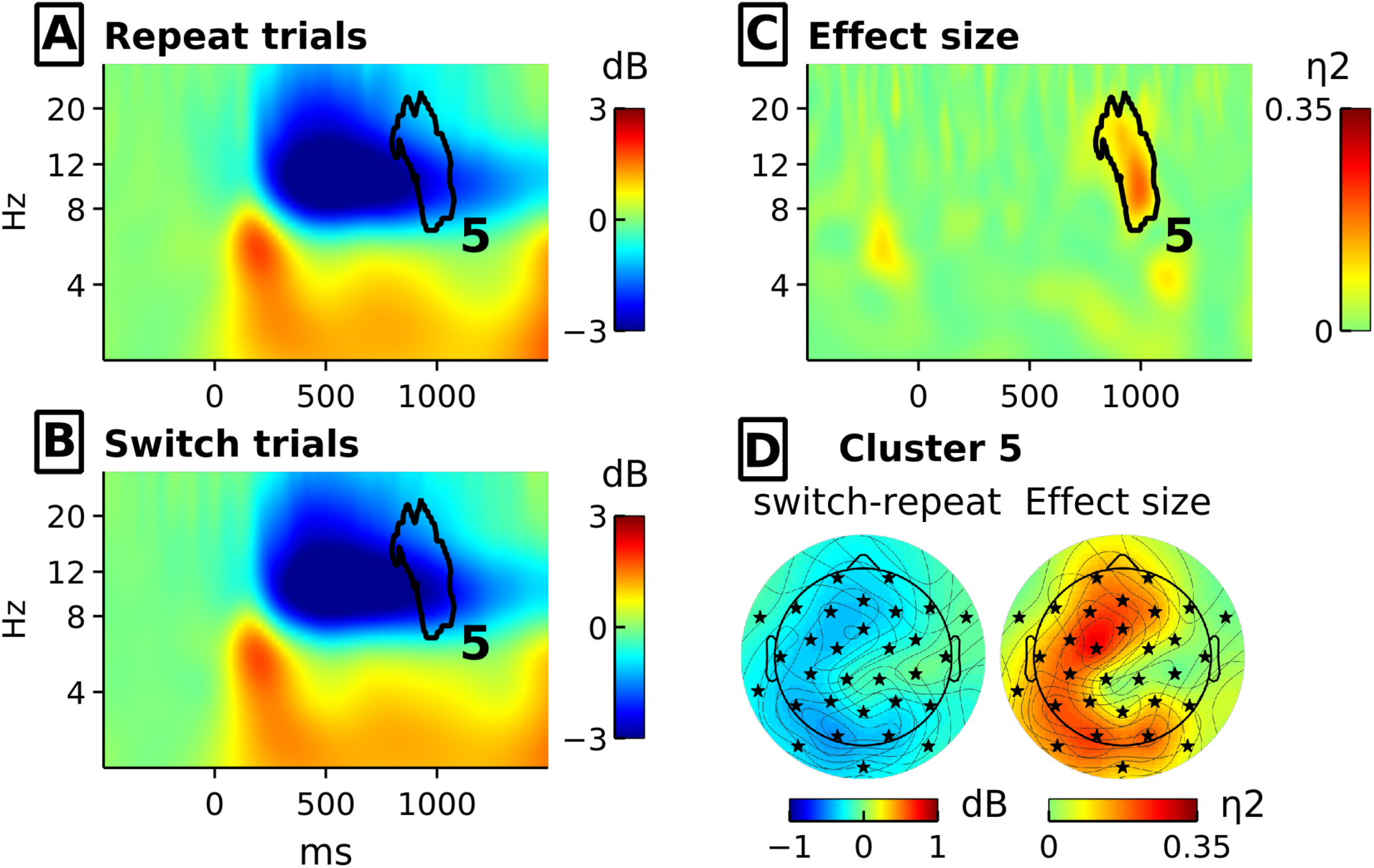
This figure illustrates the significant cluster for effects of trial sequence. The ERSPs averaged across all channels for repeat trials are depicted at [A] and for switch trials at [B]. [C] illustrates the effect sizes. The black contour lines indicate significant clusters and the black number 5 represents the cluster number. Panel [D] depicts the topographies for Cluster 5, the power difference between repeat and switch trials is illustrated at the left side and the corresponding effect sizes are illustrated at the right side. The black asterisks indicate channels with significant effects of Cluster 5.

## Discussion

In the present study we intended to obtain an electrophysiological signature of mind wandering. To this end we compared mind wandering episodes to on-task episodes in time × frequency × sensor space of the EEG. Details of the experimental design, signal processing parameters, and statistical methods were chosen with respect to avoiding as much a priori assumptions as possible. The behavioral data clearly indicate the validity of the task-switching paradigm and the probe-caught method to assess mind-wandering. Participants responded significantly slower to the imperative stimulus in switch as compared to repeat trials, which was a very large effect (η^2^= 0.73). In contrast, no differences between both trial types could be observed in terms of accuracy. The opposite pattern emerged from the comparison of mind wandering to on-task episodes. Accuracy was significantly reduced in mind wandering trials, which was a medium to large effect (η^2^ = 0.19), but no effect was visible in response times.

The largest effect of mind wandering in the EEG data, with respect to its extension in time × frequency × sensor space as well as with respect to the associated statistical effect sizes, was an increased alpha power in the inter trial interval in mind wandering episodes (Cluster 1 and 4, depicted in Figure 3). This effect was significant at all electrodes and largest over the central and parietal cortex, where it gains large effect sizes (up to η^2^ = 0.21). This finding is consistent with previous research on electrophysiological correlates of mind wandering that used the probe-caught method to assess mind wandering. A significantly increased alpha power prior to probe-caught reports of mind wandering was observed by Baldwin and colleagues (2017) during simulated driving and a vigilance task. Baird et al. (2014) found a spatially distributed effect of mind wandering in terms of a larger decrease of alpha power in response to a stimulus in mind wandering episodes during a vigilance task. Since they applied a condition-specific baseline in order to calculate the ERSPs, it might be the case that this effect actually reflects a larger alpha power in the baseline period, that is the inter-trial interval. A condition specific baseline forces the baseline to be zero and the higher baseline-power is then reflected by larger relative values in post stimulus onset episodes.

An increased alpha power in non-stimulus locked segments of the EEG is a well-known phenomenon in the research on mental fatigue (Arnau et al., 2017; Fan et al., 2015; Getzmann et al., 2018). In the context of prolonged cognitive performance, alpha power was found to increase not just as a function of time on task (Arnau et al., 2017; Fan et al., 2015; Getzmann et al., 2018), but also as a function of task load (Getzmann et al., 2018; Wascher et al., 2018). These findings mirror recent research on mind wandering frequency, which was also found to be increased when a task is well learned and little demanding (Baird et al., 2012; Cunningham et al., 2000; Giambra, 1995; Smallwood et al., 2009). It has been discussed that mind wandering might play a crucial role in the time-on-task-related decline in performance in experiments on mental fatigue (Pattyn et al., 2008). Additionally, a higher alpha power in posterior brain areas has been linked to an internally oriented cognitive state (Hanslmayr et al., 2011). Taken together, the increased alpha power in the inter-trial interval observed in the present study can be interpreted as a correlate of a redirection of attentional resources away from sensory input towards internal processing that is experienced as mind wandering. As a side note, this notion would also support the idea of mind wandering being at least partially causal to the performance decrement observed in mentally fatigued individuals. In the present study, the likelihood for mind wandering increased with time on task (see Figure 5). At the end of the experiment, however, mind wandering likelihood dropped again, probably due to anticipating the end of the experiment.

**Figure 5:**
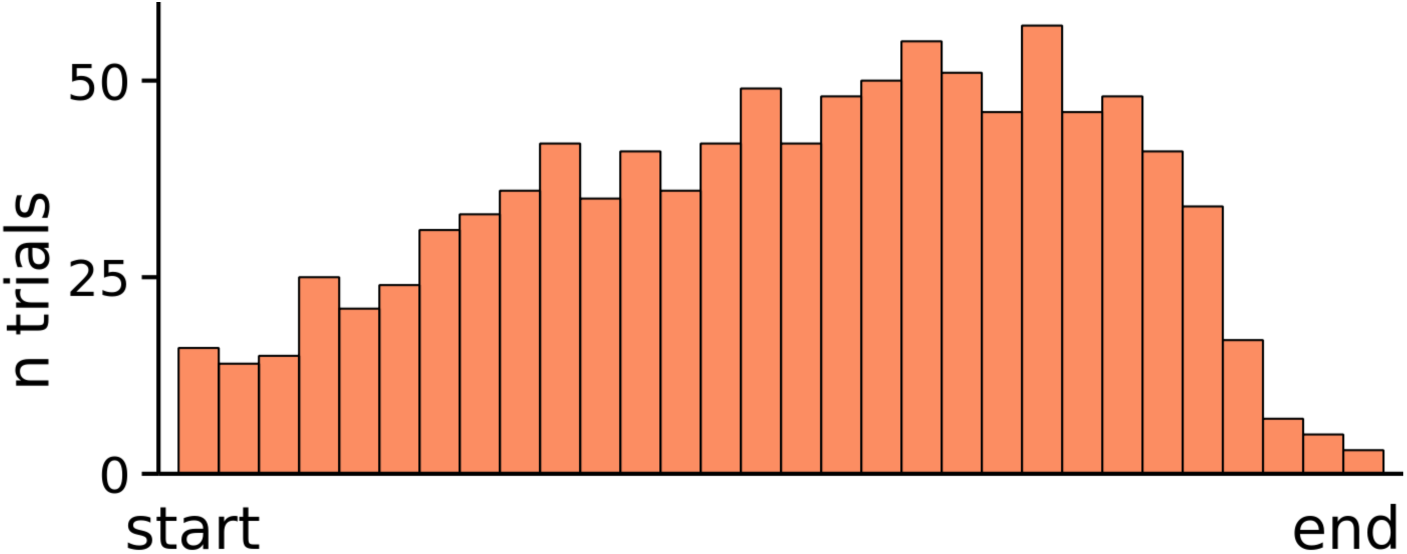
This figure depicts the distribution of reported mind wandering cumulated across all participants of the study over the course of the experiment.

The observation of an increased alpha power in mind wandering episodes in the inter-trial interval is not in line with findings of studies on mind wandering using the self-caught assessment method, which found lower alpha power in episodes prior to the button press indicating self-detected mind wandering (e.g. Braboszcz & Delorme, 2011; van Son et al., 2019). It might be the case that these episodes identified via the self-caught assessment of mind wandering rather reflect metacognitive awareness (Smallwood & Schooler, 2006). As a consequence, the spectral properties of these episodes might differ from those of mind-wandering episodes as well.

A further effect of mind wandering in time × frequency × sensor space of the EEG observed in the data is an increase of power in the lower alpha and theta range during the inter-stimulus-interval between the offset of the fixation cross and the onset of the imperative stimulus (Cluster 2 and 3). An increased theta power in response to an informative stimulus has been linked to the allocation of attentional resources (Cavanagh et al., 2012; Cavanagh & Frank, 2014). The fixation cross can certainly be interpreted as an informative stimulus that even bears the potential of disrupting mind wandering. The observed effect thus might be interpreted as a compensatory recruitment of additional resources when getting aware of mind wandering. However, theta responses as a correlate of cognitive control are usually located over the frontal cortex (e.g. Onton et al., 2005). Since the spatial distribution of the present finding is rather indistinct, the interpretation of an increased theta as a counteractive measure to compensate mind wandering remains speculative.

Mind wandering did not seem to affect task processing per se, as no significant difference between mind wandering and on-task episodes in oscillatory dynamics in response to the imperative stimulus could be observed. Other studies (e.g. Barron et al., 2011; Kam & Handy, 2013) found that mind wandering affected task processing as reflected by changes in the ERP. Barron et al. (2011), however, did not use a cue before stimulus onset, which may explain why they observed effects of mind wandering in the post imperative stimulus period that we observed in the pre-imperative stimulus period. Furthermore, there were no significant interaction effects between the factors mind wandering and trial type, that is switch versus repeat trials. This is in so far in line with previous research, as Barron and coworkers (2011) observed not only reduced P3b amplitudes to target stimuli, but also reduced P3a amplitudes in response to standard stimuli when mind wandering. They concluded that mind wandering goes along with a general suppression of external stimuli rather that being a state of suppressed central executive functioning or a state of distraction.

Although the methods used in the analysis were chosen to deliver a clear picture of the representation of mind wandering in the EEG, there are also some inherent disadvantages. This becomes obvious when considering that no effects in the theta range at frontal areas were significant for the comparison of switch versus repeat trials, although this is a common finding for this kind of task (e.g. Cunillera et al., 2012). This comparison has not been discussed here, but it shows that it might be hard to detect smaller, in the sense of spatially less distributed, effects with a cluster-based approach that comprises all channels. It thus might be the case that a more hypothesis-driven approach focusing on frontal electrodes would have been more appropriate for the purpose of detecting mind-wandering-related differences in executive functioning.

Overall, the findings of the present study clearly show that an increase of spectral power in the alpha band constitute an electrophysiological correlate of mind wandering. In combination with recent findings of a non-stimulus-locked increase of alpha power in the context of mental fatigue, this might be interpreted as further evidence for the association of a spatially distributed increase in alpha power and a bias of attentional resources towards internal processing (c.f. Hanslmayr et al., 2011). The alpha effect was located primarily in the inter-trial-interval. On the one hand, this has methodological implications as future research on mind wandering should account for this by choosing appropriate parameters for analysis. A baseline normalization procedure, for example, should allow for observing mind-wandering related variance in the baseline period.

There are also theoretical implications of the alpha effect being present specifically in the inter-trial-interval. The fact that mind wandering does not occur randomly, but instead when it is unlikely to be detrimental to performance, might indicate that it is adaptive behavior (c.f. Mooneyham & Schooler, 2013). Recent studies identified planning (Baumeister & Masicampo, 2010; D’Argembeau et al., 2011; Smallwood et al., 2009) and the fulfillment of intentions (Rummel et al., 2017; Seli et al., 2018b; Steindorf & Rummel, 2017) as potentially adaptive functionalities of mind wandering. According to a recent framework of Kurzban et al. (2013), the human cognitive resource management system will reallocate cognitive resources that are not absolutely necessary for adequate performance in a given task to another task in order to maximize the combined utility. Many tasks, however, simply cannot be engaged in simultaneously. In this context, mind wandering might be conceptualized as a collective term for all the tasks an individual is able to engage in while performing another task, which could explain its high prevalence (Kane et al., 2017; Killingsworth & Gilbert, 2010). The downside of such an optimization strategy is that unexpected and critical situations may lead to primary task failure (e.g. Galéra et al., 2012; Qu et al., 2015).

Finally, the identification of alpha power as a correlate of mind wandering may also have practical implications. It should be evaluated whether the online detection of alpha-power-increases in working environments prone to mind wandering is feasible in order to detect mind wandering and prevent performance failures.

## Funding

This work was supported by the Excellence Initiative of the German Research Foundation (DFG) [grant number ZUK 49/Ü 5.2.178].

### Acknowledgements

We would like to thank Emily Brett, Isabel Gebhardt, Sarah Hladik, and Niklas Neumann for their assistance in collecting the data as well as Gidon Frischkorn for programming the experiment.

